# Avian Influenza in Ireland: A Spatiotemporal, Subtype, and Host-Based Analysis (1983-2024)

**DOI:** 10.1101/2025.05.26.656175

**Authors:** Maeve Louise Farrell, Guy McGrath, Laura Garza Cuartero, Gerald Barry

## Abstract

Avian influenza virus (AIV) is a significant global concern, causing widespread mortality in wild birds, domestic poultry and most recently wild and domestic mammals. This study presents a retrospective analysis of AIV detections in the Republic of Ireland. Data was sourced from official surveillance databases, peer-reviewed literature and grey literature sources. Spatio-temporal, host-specific and subtype patterns were assessed using descriptive statistics, chi-square tests, linear regression and kernel density estimations. A total of 2,888 confirmed AIV detections were recorded from 25 of Ireland’s 26 counties. Wild birds accounted for 98.7% of detections, with domestic birds comprising 1.3% and two detections in foxes. H5N1 was the most prevalent subtype (96.7%) followed by H5N8 and H6N1. Spatial clustering was observed in urban areas, particularly Dublin. The highest seasonal peak occurred during summer, contrasting with traditional winter-associated patterns. Several detections occurred in migratory species outside of typical residency periods, suggesting potential climate-related shifts in migration behaviour. This study represents the first review of AIV surveillance data in Ireland to date. The findings highlight evolving patterns in virus distribution, seasonality and host dynamics, with implications for national surveillance strategies. Continued cross-species monitoring and integration of ecological data are essential to inform effective management strategies.

## 1. Introduction

Avian influenza virus (AIV) is a global threat to both human and animal health, with high levels of morbidity and mortality (Charostad et al., 2023). Belonging to the *Orthomyxovirdae* family, these viruses are classified as influenza A, and subtyped based on two major antigens, haemagglutinin (HA) and neuraminidase (NA), (Tan., 2012). These viruses are further categorised by their pathogenicity; low pathogenic avian influenza (LPAI) and high pathogenic avian influenza (HPAI) based on their ability to infect poultry (European Food Safety Authority, 2025). While LPAI, as the name suggests, has milder symptoms, HPAI - particularly the global H5N1- can cause outbreaks of disease, with high mortality rates in wild and domestic birds (Department of Agriculture, Food and the Marine., 2024).

Wild avifauna, particularly waterfowl and other water-associated birds, are recognised as key reservoirs of both pathotypes (Bowedes and Kuiken, 2018). These species, alongside migratory birds, play a central role in the global dissemination of AIV through seasonal migrations, often disseminating the virus across continents (Kandeil et al., 2023). In recent years, spillover events have involved mammals, such as mesocarnivores (foxes, seals) and more recently livestock, raising concerns about the expanding host range and potential for zoonotic transmission (Peacock et al., 2025). With the recent outbreak of HPAI H5N1 2.3.4.4b in American cattle, cases have been reported in humans exposed to these animals, most commonly workers on farms (Mostafa et al., 2024).

In 2022 alone, HPAI outbreaks led to the culling of an estimated 50 million birds in affected European countries (Adlhoch, 2023), causing not only extensive mortality, but also disruption to poultry production and economic losses. The increasing prevalence and geographic spread of such events (Parums., 2023) underscores the need for not only enhanced surveillance, but also understanding into the various factors that could be causing outbreaks.

Ireland is a nation located in Northwestern Europe, and is divided into four provinces: Leinster, Munster, Connacht, and Ulster, with 26 counties forming the Republic of Ireland. As of its most recent population census, Ireland is home to over 5 million people (Central Statistics Office, 2024), making it one of the smaller nations in terms of population within the European Union. Agriculture is a vital sector of the Irish economy, alongside services and technology. The poultry industry in Ireland, while smaller than other agricultural sectors, plays an important role. It is estimated that a significant proportion of poultry farming is commercial, with intensive operations concentrated in regions like Monaghan, Cavan, and Cork (Kelleghan et al, 2020). The rest of the poultry population is raised in semi-commercial or backyard systems, which are more prominent in rural areas across the country. Backyard systems are generally small-scale and often integrate poultry with other forms of subsistence farming, relying on both traditional and improved breeds to meet household needs. Furthermore, Ireland plays a niche role as a major flyway for migratory birds, alongside its mild climate and abundance of wetlands (Crowe et al., 2009).

Despite extensive global research on AIV, several key gaps remain at a national context. In Ireland, there is a lack of longitudinal analysis of AIV surveillance, limiting understanding of spatial, temporal and potential host-specific trends. This study aims to address this by conducting a desk-based review and analysis of AIV detections in the Republic of Ireland from 2003 - 2024. This research aims to characterise the temporal, geographical and host-based patterns of AIV occurrence, and assess the diversity and distribution of viral subtypes. The findings are intended to inform future surveillance priorities, guide national policy, and support targeted research efforts, contributing to more effective disease monitoring and control strategies within the Irish context.

## 2. Methods

### 2.1 Data Collection

A comprehensive literature search was conducted to collect data on AIV detections in the Republic of Ireland from the earliest reported case up to December 2024. The search included both peer-reviewed and grey literature sources to ensure completeness and accuracy. Primary sources consisted of official reports and databases from the Department of Agriculture, Food and Marine (DAFM), International System for Agricultural Science and Technology (AGRIS), World Animal Health Information System (WAHIS), and the European Food Safety Authority (EFSA). Additionally, academic databases such as PubMed were queried using search terms including “avian influenza” AND “Ireland.”

Retrospective molecular data from polymerase chain reaction (PCR) testing was included. Data from early detections up to December 2024 were included to provide a comprehensive temporal analysis of incidence.

### 2.2 Data Extraction and Curation

Data was imported into the web-based platform Google Sheets, with outbreaks recorded as individual rows in the dataset. All cases were georeferenced to county level and included associated metadata when available. In the case of commercial farms, approximate coordinates were used to ensure privacy and protect farm locations. Domestic animal cases were aggregated as single entries due to the absence of specific animal counts, as per national guidelines on infection management. Due to insufficient data regarding the exact number of domestic animals affected during an outbreak due to national guidelines regarding clearing of infection, it is not possible to identify exact incidence in terms of population numbers. The dataset included key variables such as species, date of detection, county, geo coordinates (if available), Influenza A RNA detection status, and subtype. Species were taxonomically classified and grouped into ecological guilds to facilitate ecological analysis.

Duplicate entries were removed, and records lacking confirmed Influenza A RNA detection were excluded. Records with partial data (e.g., missing date or county) were retained when sufficient metadata allowed for spatial or temporal analysis. To ensure data accuracy, all entries were cross-referenced with multiple sources when available. In cases of data inconsistencies, e.g. differing reports on date of occurrence, priority was given to official sources.

Temporal aggregation was performed at annual, monthly, and seasonal levels to detect patterns. Migratory status was assigned using avian profiles specific to the Republic of Ireland, based on published sources.

### 2.3 Statistical and Spatial Analysis

Descriptive statistics were applied to analyse the overall dataset to include frequency distribution across year, county, subtype and host category. Chi-square tests were utilised to assess seasonal variations in detections. To assess species-specific risk, z-scores, relative risk (RR) and 95% confidence intervals were calculated to identify species at a higher risk of AIV detection with relation to the overall dataset. Pearson’s correlation analysis was conducted to explore potential associations between AIV detections in wild birds and domestic poultry.

Linear regression analysis was utilised to examine trends in occurrence over time. Data analysis was carried out using Google Sheets and R Studio (2024, Version 12.1), while initial data cleaning and basic descriptive analysis were performed in Google Sheets. Statistical significance was set at p < 0.05. Visualisations, including time series plots and geographical maps, were generated in R Studio to illustrate trends and spatial distributions. ArcGIS Pro (Version 3.2.0) was utilised to perform kernel density estimations, and creation of geographic maps.

## 3. Results

### 3.1 Descriptive Characteristics of Dataset

A comprehensive review of the literature identified 2888 records of AIV in the Republic of Ireland between 1983 and 2024. Of these, 2861 were fully subtyped, one case was identified only to the hemagglutinin (H) type, and 25 cases remained untyped (Table 1). The majority of cases involved avian wildlife (n = 2849, 98.7%), while domestic avians comprised a small portion (n = 37, 1.3%). Two cases (0.07%) involved non-avian wildlife, both in red foxes (*Vulpes vulpes*), representing the only mammalian detections in the dataset. AIV records were documented in 25 counties of the 26 counties, with no records identified for Carlow and nineteen occurrences without county data.

**Table 1.**
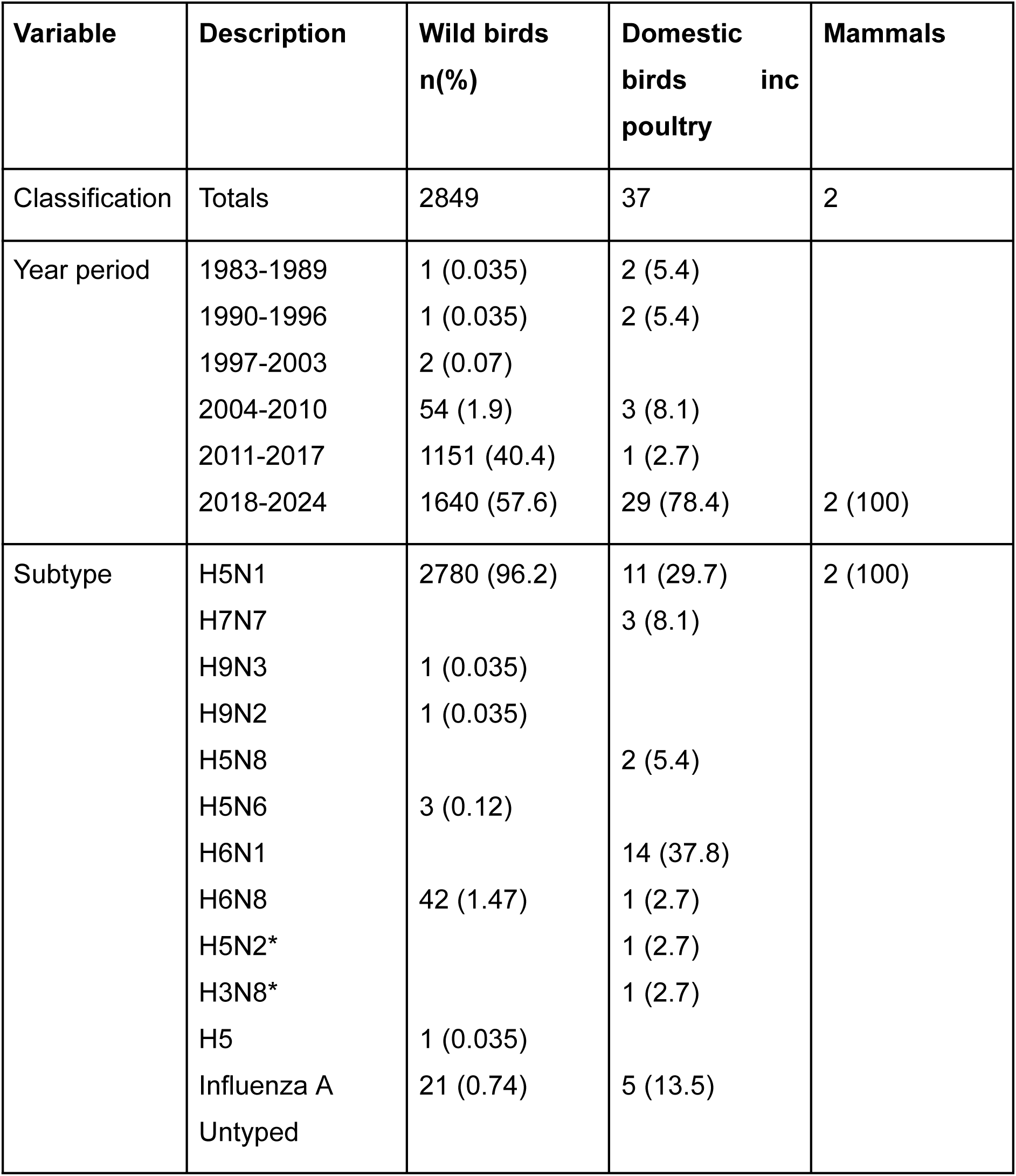
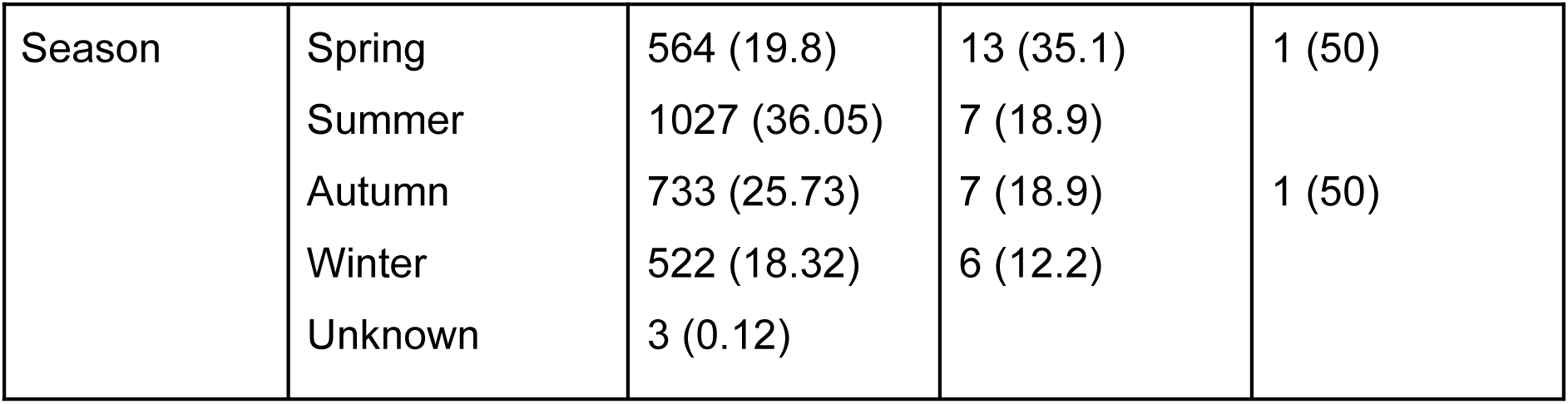
Summary of descriptive characteristics of avian influenza detections in Ireland (1983-2024)

In terms of subtype dynamics, H5N8 low pathogenic avian influenza (LPAI) was the first subtype identified in Ireland. This was followed by H7N7 and H9N3, although detections were limited. The H5N1 subtype was the most prevalent, accounting for 2,793 cases (96.7%), and was the only subtype detected continuously between 2003 and 2024. H5N6 (n = 3) was first detected in 2018, and H5N8 was observed intermittently, with detections in 1983, 2016, 2017, 2020, and 2021. H5N6 and H5N3 were each only observed in isolated years.

### 3.2 Temporal Distribution of Detections

The earliest detection of AIV in domestic avians occurred in 1983, involving HPAI H5N8 in a turkey flock and a duck (McNulty et al., 1985; Kawaoka et al., 1987). Subsequent cases appeared sporadically; once in 1989, 1993, 1997, and twice in 1995. However, detailed information from these outbreaks, such as county, or other contextual data, is not available. No further cases were reported in domestic species until a singular outbreak in 2007, followed by two outbreaks in 2009. A total of 37 cases were identified from 1983 to 2024, encompassing various domestic bird types including turkeys, ducks, chickens, laying hens, broilers, and commercial flocks.

In wild avifauna, the earliest cases were recorded in 1983, where the H5N8 and H9N3 subtype were identified in an unknown duck (*Anas spp.*) and a mallard (*Anas platyrhynchos*), respectively. In 2003, H5N1 was detected in a Mew Gull (*Larus canus*) in County Galway. Following these initial detections, no further cases were reported until 2005, when seven additional H5N1 cases were confirmed across multiple avian species. Between 2006 and 2009, annual detections remained low, ranging from two to nine cases per annum until 2010 when an increase to nineteen cases was noted.

A substantial increase in AIV detections occurred in the subsequent years, with cases ranging from 21 to 47 until a pronounced increase occurred in 2015, where 219 cases were identified. The period between 2015 and 2019 represented the highest phase of transmission, which was characterised by numerous cases (n= 1885, 65.3%), with the highest recorded case count occurring in 2019 (n = 874, 30.2%).

Seasonal patterns indicated the highest detections in August (n=439, 15.2%), with summer being the most prevalent season overall (n=1034, 35.8%), (Figure 3.1A). A total of seven cases lacked seasonal classification and were excluded from this analysis. A chi-square test indicated significant variation in detections across seasons (χ2 = 216.68, *df =* 3, p < 0.05), with summer contributing the highest individual chi-square value (χ2 = 136.67).

**Figure 3.1.**
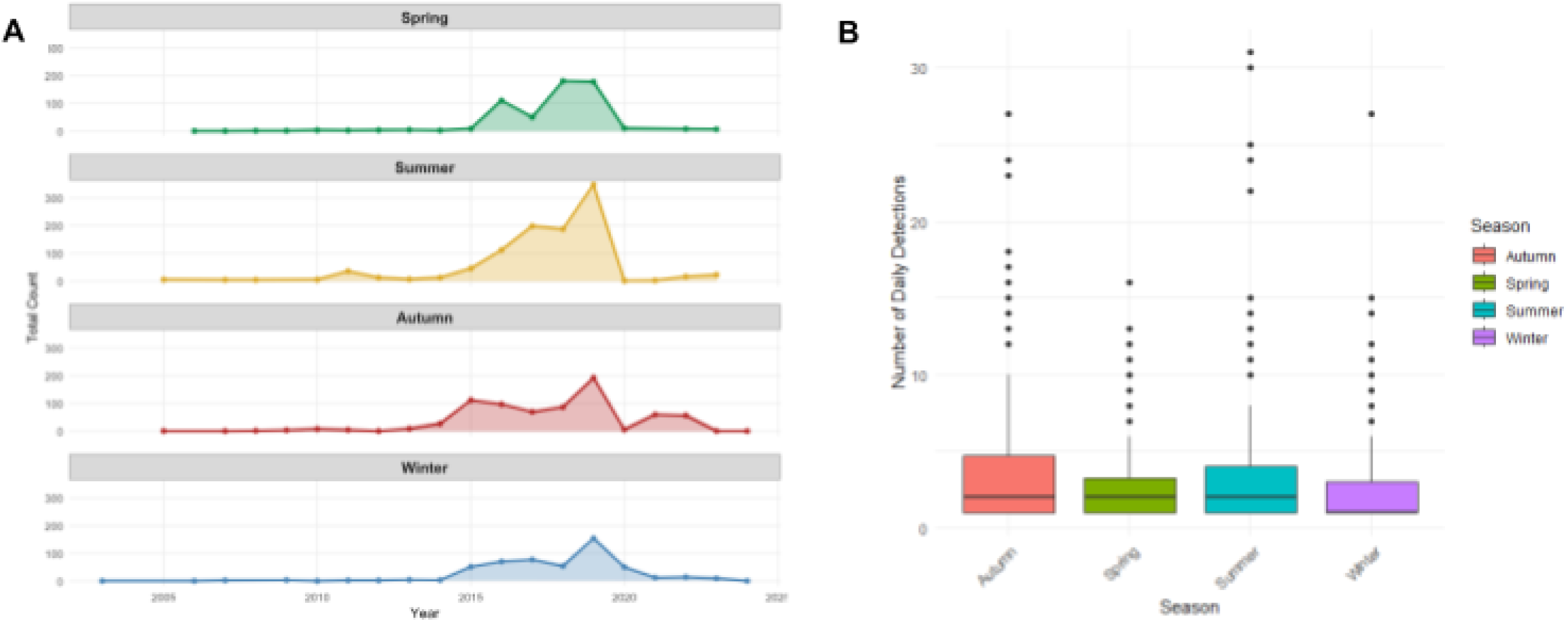
A. Faceted chart detailing temporal distribution of AIV detections across the four seasons; Spring, Summer, Autumn and Winter. B. Distribution daily AIV detections by season.

Furthermore, all seasons had significant day-to-day variability in detections. Autumn had the highest median daily detection rate (3.82; range: 2–17), followed closely by summer (3.59; range: 2–31), as illustrated in Figure 3.1B, indicating seasonal concentration and within season fluctuation in AIV detections.

### 3.3 Geographic Distribution of Cases

Geospatial analysis identified County Dublin as the county with the highest AIV detections (n = 1,197; 41.45%), followed by Galway (n = 411; 14.23%) and Cork (n = 212; 7.34%). Data prior to 2003 lacked location details and was therefore excluded. Notable geographic and temporal expansion were observed over the years. As illustrated in Figure 3.2, early detections of AIV (2003-2010) in wild avifauna were relatively sparse and largely confined to coastal regions. During the years that followed (2011-2017), detections became more widespread, with emerging clusters in eastern counties, such as Dublin and surrounding areas. In the most recent years (2018-2024), AIV detections were reported across nearly all counties, with marked increase in density in the east, alongside substantial clustering in parts of the midlands and west.

**Figure 3.2.**
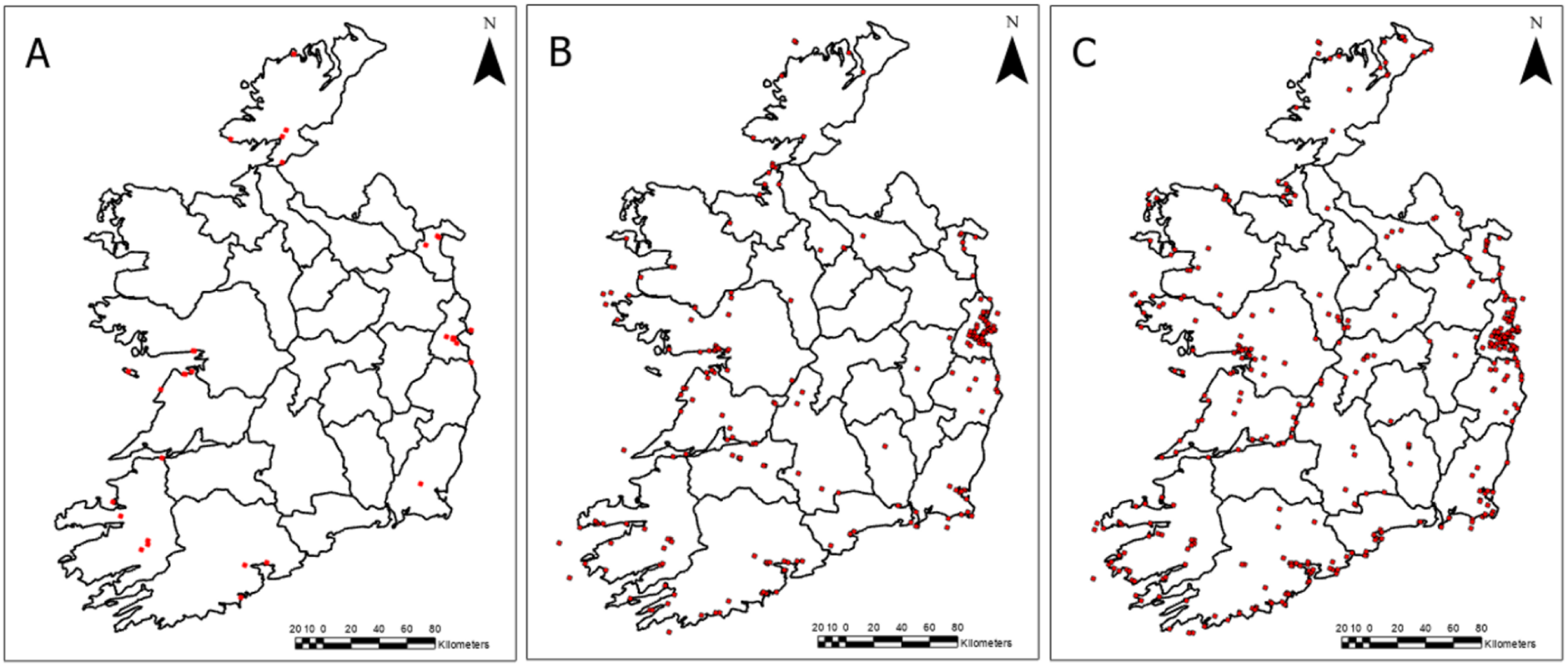
Spatiotemporal distribution of AIV detentions in the Republic of Ireland (2003-2024).

Distribution of the dominant subtype, H5N1, was variable across the counties, reflecting broad geographic spread. In contrast, other subtypes were more spatially restricted. H5N2 was detected in Tipperary and Wicklow, however the latter was in combination with H3N8 in a domestic flock. H5N3 was detected in Louth (n=2), while H5N6 was detected in both Tipperary (n=2) and Clare(n=1). H5N8 demonstrated the largest spatial range among non-H5N1 subtypes, with detections in fourteen counties, namely Cork (n=10), Galway (n=5), and Monaghan (n=4), with two cases occurring in unknown counties.

A kernel density estimation was performed on avifauna data using ArcGIS Pro to visualise the spatial distribution of AIV across Ireland from 2003-2024. As seen by Figure 3.3, spatial heterogeneity in distribution of AIV detections was seen. Density values ranged from 0.001 to 10.112 cases per km^2^, with highest densities seen in Dublin. Most clusters of density occurred in coastal regions, such as in Cork and Galway. Lower density occurrences were dispersed throughout midland and western counties.

**Figure 3.3.**
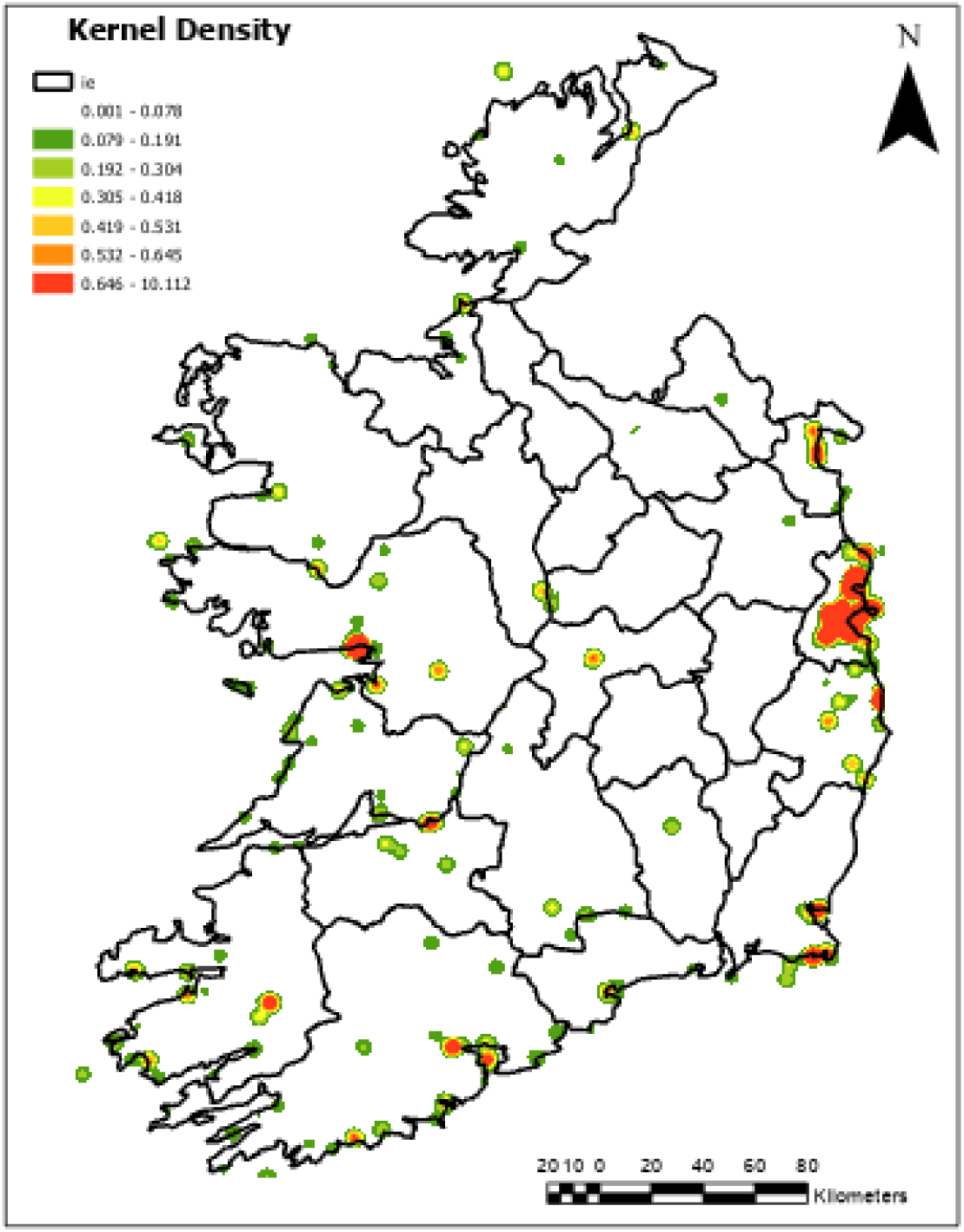
Kernel density estimation of the geographical distribution of avian influenza virus (AIV) detections in the Republic of Ireland from 2003 to 2024. Density was visualised using standard deviation (SD) contours, with intervals set at 1 SD. A red-to-green color scale was used to indicate relative density, with red representing areas of higher detection concentration and green indicating lower density.

In contrast, density of detections in domestic birds were more geographically confined, occurring primarily in counties with established poultry production. Notably, Monaghan emerged as a key hotspot, consistent with intensive commercial poultry operations. Additional outbreaks were identified in Wicklow, Cavan, Tipperary and Kerry, as seen in Figure 3.4.

**Figure 3.4.**
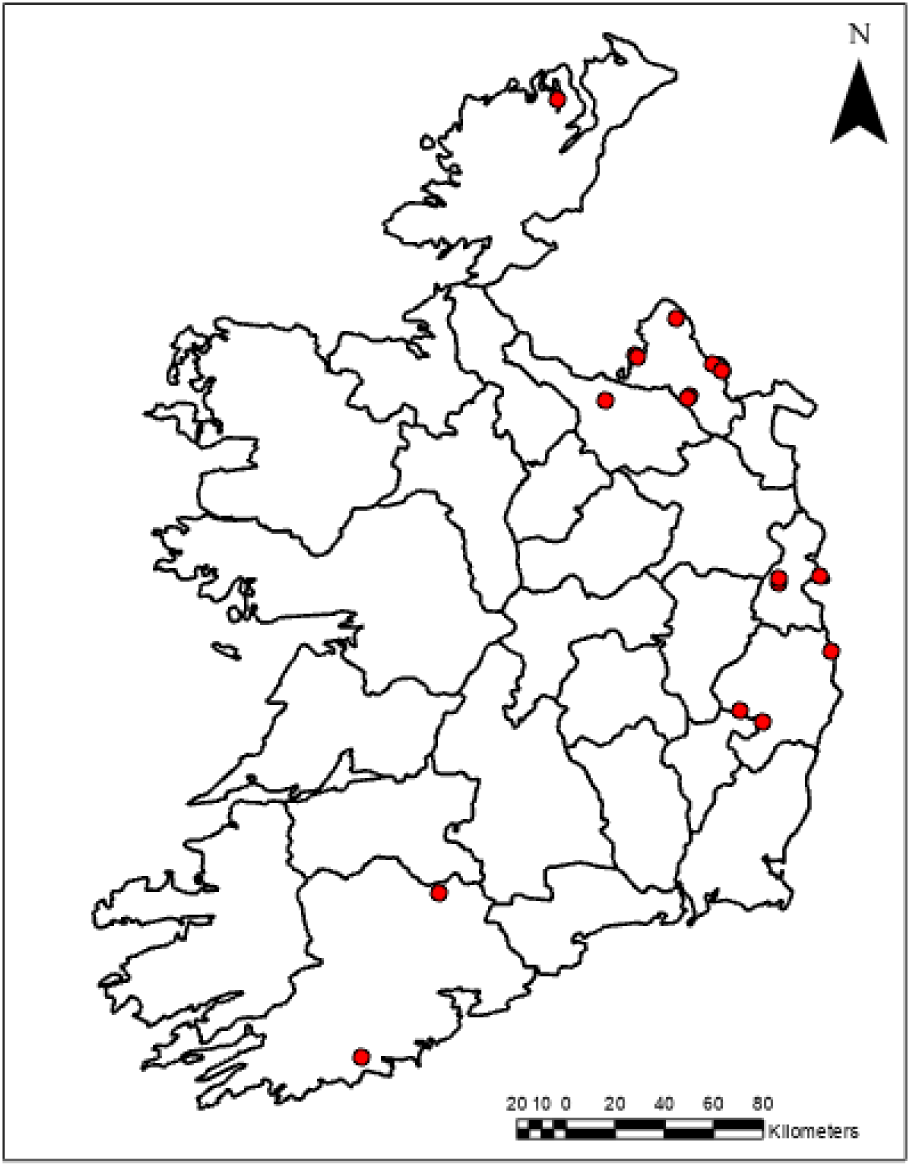
Spatial distribution of AIV detections in domestic birds in the ROI (2003-2024).

### 3.4 Host-Based Patterns

Overall, 39 genus were identified in the dataset, with 28 entries unable to be categorised to a certain genus (namely domestic avians with no species listed). Analysis revealed distinct trends in the occurrence of different genera, with significant variances seen across the datasets. The most frequently affected genus was *Cygnus* spp. (n=454, 15.72%), *Chroicoceohalus* spp. (n=340, 11.8%), *Ardea* spp. (n=261, 9.04%), *Larus* spp. (n=231, 8%) and *Egretta* spp. (n=227, 7.86%). Among these, *Cygnus* spp. had the most overall detections with three different subtypes detected; H5N1 (n=416), H5N8 (n=32), and H5 (n=1). Z-scores were calculated to identify genus with higher relative risk for AIV detection. Furthermore, the relative risk score for *Cygnus* was 5.97, with a z-score of 3.5, indicating significant increases in detection relative to other genera. *Chroicoceohalus* had a z-score 2.44 and RR 4.47, whilst *Ardea* (z-score 1.56, RR 8.5. All exhibited significantly elevated risk, however these genus contributed substantially to the overall cases, with subsequent genera showing gradual decline in frequency.

At a species level, Black-headed Gull (*Chroicocephalus ridibundus*) emerged as the most frequently documented species (n=340, 11.77%, z 3.67, RR 6.66). *Cygnus olor* (Mute Swan) represented the second most prevalent species (n = 300, 10.39%, 95% CI [9.28%, 11.50%]), with a z-score of 3.16 and relative risk of 5.87. This was followed by *Ardea cinerea* (Grey Heron) with 261 observations (9.04%, 95% CI [7.99%, 10.09%], z-score = 2.66, RR = 5.11). Subsequent species had gradual declines in frequency, whilst maintaining statistically significant deviations from the mean, e.g. Little Egret (n=227, 7.86%, z 2.23, RR 4.44). The z-scores indicate significant non-random distribution of species occurrence, with the three species with most occurrences (*C. ridibundus, C. olor, and A. cinerea*) having >2.5 z-scores, suggesting an overrepresentation in surveillance data. Eleven species had over 100 cases of AIV, whilst 23 species had less than five detections.

The majority of detections (n=1299, 44.98%) occurred in waterfowl, followed by seabirds (n=657, 22.75%), waders (n=492, 17.04%) and raptors (n=244, 8.45%). Migratory species accounted for the majority of detections (n=1574), with species considered partially migratory accounting for 296 detections. The first detection of AIV in wild birds occurred in 2003 occurred in a migratory species, however during the next detections in 2005, four out of the seven detections occurred in non-migratory species. With the exception of 2007, 2008, 2010 and 2024, detections were higher in migratory species compared to non-migratory species.

A Pearson’s correlation analysis was conducted using monthly aggregated data to assess the potential for a relationship between AIV in wild avifauna and domestic birds. The correlation coefficient was −0.75 (p = 0.7167), indicating no statistically significant linear relationship.

### 3.5. Subtype Distribution

Between 1983 and 2024, nine AIV subtypes were detected in Ireland, with 26 records untyped and one typed as H5Nx. H5N1, first recorded in 2003, predominated the distribution (n=2793), followed by H5N8 which was detected first in 1983. In contrast, H6N1, and H5N6 were detected fourteen, and three times respectively, both occurring in singular separate years. Considerable variability in subtype distribution was observed across the years, with a median of 70.44 cases per year.

H5N1 occurred year round, consistent with its endemic presence during the study period. By contrast, H5N6 had singular detections in January, February and March. H5N8 had a strong seasonal pattern, with the highest frequency in December (n=20), followed by January (n=9), February (n=7), November (n=7) and October (n=1) as seen in Figure 3.5.

**Figure 3.5.**
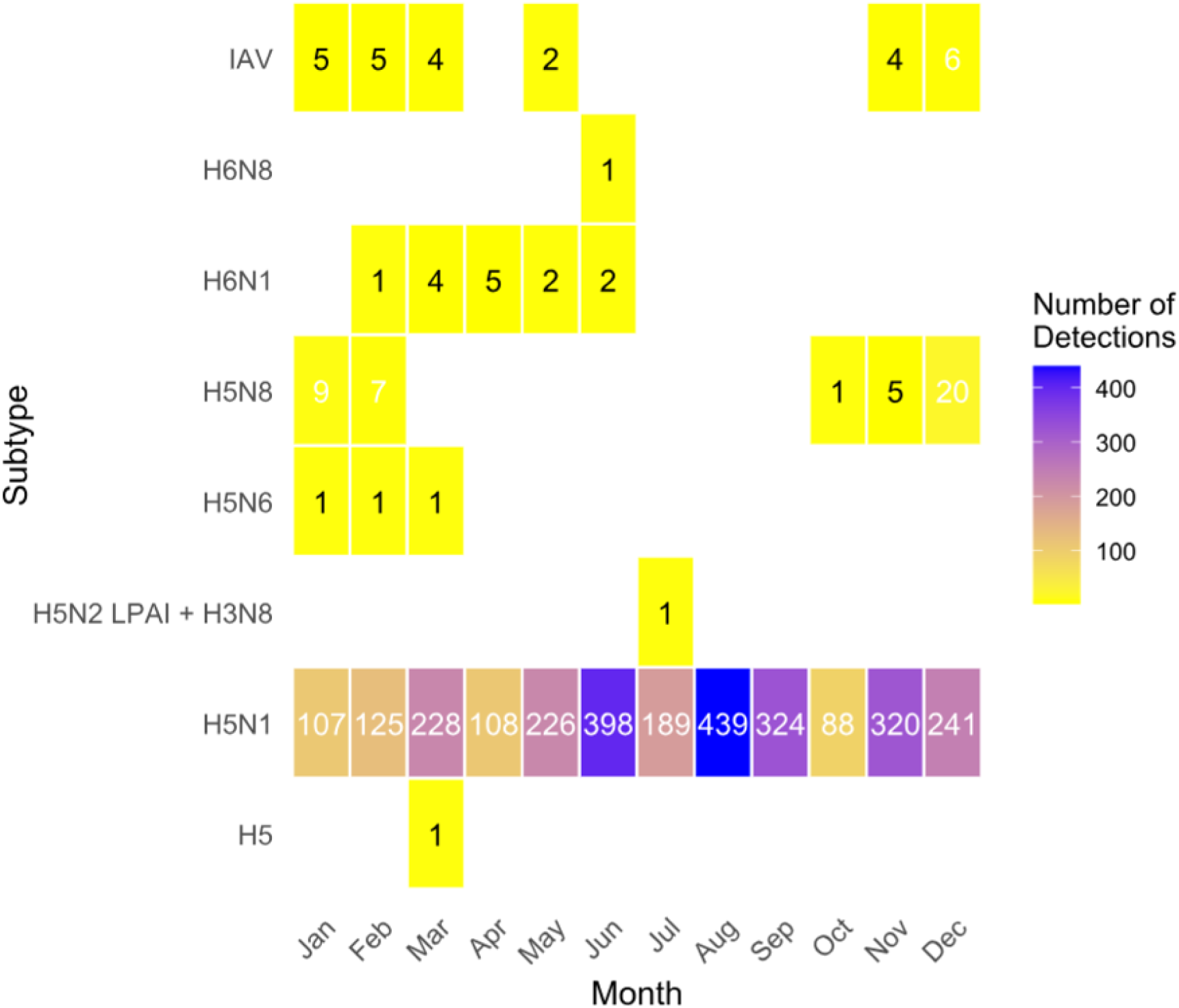
Seasonal distribution heatmap of avian influenza virus (AIV) detections by subtype.

Among the domestic bird cases, H5N8 and H5N1 were the most frequently detected subtypes. H6N1 LPAI was also prevalent, particularly in laying hen flocks in Monaghan during 2020. The co-occurrence of multiple subtypes was observed in a duck game flock in Wicklow in 2021 (H5N2 LPAI and H3N8).

The year 2021 demonstrated the greatest diversity in terms of subtype detection, with concurrent detections of H5N1, H5N8, and H6N8, and a co-occurence of H5N2 and H3N8 occurrences in a duck flock, as seen in Figure 3.3. Following linear regression analyses, H5N1 showed slight positive changes over time, however this was not statistically significant (p=0.153). Considering overall detections, a regression model indicates a slope of approximately 7.6 detections/year (p=0.044), suggesting a slight upward trend over time.

## 4. Discussion

This research presents a comprehensive analysis of 2,888 records of AIV detections in the Republic of Ireland over a 41 year period. The substantial increase in reported detections, in particular between 2015-2019, may be indicative of either a rise in prevalence or an increase in national surveillance efforts. A notable finding of this research is the seasonal distribution of detection, which peaked during the summer months, in contrast to the common viewpoint that AIV is predominantly a winter-associated pathogen (Ahmed et al., 2023, Tuncer et al., 2013). This contrast warrants further investigation into potential ecological drivers and host behaviours that could influence virus transmission.

The findings of this study are consistent with global patterns identifying wild avifauna as reservoirs of AIV (Ren et al., 2025; Yang and Fan, 2025). Occasional detections of AIV in mammals may reflect environmental contamination, or potentially scavenger-mediated transmission pathways. These observations highlight the need for further research into the role of non-avians in prevalence of AIV (Peacock et al., 2025; Velkers et al., 2017). To the best of the authors knowledge, this is the first study to systematically collate and analyse AIV records in Ireland, spanning over four decades of data.

Beyond surveillance related explanations, It is essential to consider broader ecological and behavioural factors that could influence temporal increases in detection. For example, while increased surveillance (Department of Agriculture, Food and the Marine., 2024), and fluctuations in migratory bird populations are likely contributors, long-term data indicates significant shifts in bird populations. A report issued in 2019 detailed that wintering waterbirds suffered a decrease of 40% in the previous 17 years. However, some species, such as the Whooper Swan, saw increases in recent years, with a 13.4% change (Burke et al., 2018). These alterations in host dynamics suggest the possibility of altered migratory pathways in Ireland, potentially influenced by climate change. For example, detections of AIV in birds outside of ther normal period of residency was seen in this research. Notably, Whooper Swans (n=151), a species that mainly overwinters in Ireland October to April in Ireland, had 19 detections occurring outside of this period. Similarly, the Northern Pintail (n=40), a winter visitor, had two detections in June, while the Greater Scaup (n=37), had three recorded detections in May. The Common Eider (n=44), another wintering species, had ten detections outside of its expected seasonal range, as seen in Supplementary Material. These irregularities could reflect changes in migration patterns or shifts in overwintering timepoints, potentially driven by climate change (Rubolini et al., 2010), which may also be impacting virus dynamics, evolution and transmission (Prosser et al., 2023).

Species-specific trends reinforce the importance of surveillance and research into certain hosts and host dynamics. Increases in detection over time in species such as swans (*Cygnus* spp.) align with existing literature recognising them as a key reservoir of both low and high pathogenicity (Bergervoet et al., 2019; Lambrecht et al., 2016). The significantly increased z-scores for these birds indicate non-random patterns of infection, potentially driven by habitat based exposures. Water-associated birds such as swans, gulls, herons and egrets are well-documented reservoirs of AIV due to their presence in aquatic environments (Sheikh et al., 2025; Soda et al., 2022; Woo et al., 2017)

Spatial clustering of AIV detections in Ireland appear to be influenced by a combination of anthropogenic and ecological factors, including population density, proximity to water bodies and distribution of targeted surveillance strategies. The country’s capital, Dublin, accounted for approximately 41% of reported detections, a disproportionately high figure. Multiple factors could be considered drivers of this, including human population density. Results from the most recent Irish Census in 2022 revealed that Dublin accounted for approximately 40% of the total Irish population (Central Statistics Office, 2022). Furthermore, this increased population density, and the fact the central laboratories of DAFM are situated on the Dublin border, could impact reporting. In January 2021, a virtual reporting platform was created by DAFM to facilitate the reporting of birds potentially affected by AIV. To date, 3519 reports have been submitted via this platform (January 2021- March 2025), with highest reports submitted from areas of dense human population and coastal counties (Department of Agriculture, Food and the Marine, 2025). Kernel density estimations confirmed these findings, with high density clusters visualised in Dublin, Galway and Cork regions.

As seen by Figure 3.1 in the Avian Check App report, a spike in reports was visualised in September and October of 2022, coinciding with increased AIV cases in Gannets globally (Lane et al., 2023). As seen in the data presented in this research, of the 97 detections in 2022, 52 were detected September and October, of which 48 were Northern Gannets. Similarly, in the data published by Paradell et al (2023), increased cases of HPAI were observed in gannets and an estimated minimum local population mortality of 3126 birds (95% confidence intervals 2993-3260).

The multi-county occurrence of H5N1 reflects its well-documented virulence and ability for fast transmission (Banyard et al., 2024; Mao et al., 2024). Conversely, the limited spatial distribution of other subtypes such as H5N3 and H5N6 suggests the potential for isolated introductions or localised spillover events. Interestingly, H6N1 was only detected in domestic birds in 2020, and was restricted to the county of Monaghan. This suggests potential farm to farm transmission, with additional detections occurring in Northern Ireland during the same timeframe (McMenamy et al., 2023). Whilst a wild bird source was not definable in this outbreak, it cannot be ignored that this subtype likely had a wild bird origin due to its prevalence in wild birds. Furthermore, the environment is another potential route of transmission due to the ability of LPAI to persist (Rohani et al., 2009). In the case of H5N8, this subtype was detected in a turkey flock in Wicklow in early December, with the same subtype detected in a Whooper Swan just prior. However, a distance of just under 40 km was found between these two detections. Two additional cases in swans were detected shortly after the initial case in the Whooper swan, with these three cases clustering together. The lack of statistically significant correlation between wild avifauna and domestic AIV cases in this study suggests limited direct spillover in an Irish context. These findings highlight the importance of integrating spatial and ecological data to inform surveillance strategies, and improve understanding of environmental and anthropogenic factors influence on viral distribution and persistence.

The detection of AIV in red foxes is particularly concerning as it underscores the potential for the virus to spill over into non-avian hosts. A recent Irish serological study conducted between 2022 and 2024 detected influenza A antibodies in mesocarnivore species; red foxes, Eurasian Badgers (*Meles meles*) and American Mink (*Neogale vision*). This study utilised a commercially available enzyme linked immunosorbent antibody assays (ELISA) screening 219 samples, of which 31 tested positive for influenza A antibodies. Of these, 23 of 28 antibody-positive fox samples were typed as H5, along with the singular badger sample. Two antibody-positive samples from American Mink could not be subtyped, testing negative for both H5 and H7 (Ruy et al., 2025). These findings are consistent with broader European surveillance efforts, where seroprevalence of various influenza A antibodies has been documented in multiple wild mammal species, including seals, foxes, martens, polecats, badgers, and stoats (Chestakova et al., 2023; Gholipour et al., 2017; Bordes et al., 2015). The ability of these animals to survive infection and retain antibodies suggests that wild mammals could act as incidental hosts or reservoirs of influenza A viruses, highlighting the importance of incorporating mammalian wildlife into AIV surveillance networks.

Alongside this, growing concerns for livestock have arisen given the recent spillover of HPAI H5N1 2.3.4.4b into livestock species. This follows the detection of the virus in dairy cattle across several states in the United States, confirming the first cases of HPAI in bovine populations (Baker et al., 2025). Moreover, the detection of H5N1 in a sheep in Yorkshire, United Kingdom (Mahase., 2025) further highlights the expanding host range of the virus. To date, AIV has not been detected in livestock in Ireland, nevertheless these events signify a concerning shift in the epidemiology of HPAI. The evolution of host dynamics highlights the need for enhanced cross-species and cross-sectoral surveillance, and use of One Health approaches to monitor and mitigate potential risks.

The retrospective nature of this analysis, and reliance on existing surveillance records, introduces inherent biases including reporting practices and variability in sampling efforts. Poultry data, in particular, may be incomplete due to the need to protect farmers’ privacy. Additionally, changes in surveillance intensity over time may have skewed detection rates and temporal trends. Mammalian data were both limited in frequency and geographic scope; however they highlight the need for inclusion of non-avian species in future surveillance frameworks. Despite these limitations, the findings of this analysis provide valuable insights into long-term trends and emerging risks associated with AIV in Ireland.

## 5. Conclusion

This study provides the most comprehensive longitudinal analysis of AIV in the Republic of Ireland to date, spanning across four decades and including both avian and non-avian hosts. These findings highlight clear temporal and spatial trends, with urbanised clustering. Seasonal peaks were identified during summer months in contrast to normal misconceptions that AIV is predominately a winter-associated pathogen, suggesting the influence of shifting migratory behaviours or impacts of climate change. Specific genera were more commonly affected, underscoring the importance of species-specific targeting in surveillance and monitoring efforts. This research makes important contributions to the evidence base of understanding the environmental, host and temporal related factors impacting AIV occurrence in Ireland. Further research is needed to elucidate the potential for farm-level outbreaks, the persistence of AIV in wild bird populations throughout the year, and the potential role of mammals in the dissemination of the virus.

## Acknowledgements

This project is co-funded by the European Union (Project ID: 101132970). Views and opinions expressed are however those of the author(s) only and do not necessarily reflect those of the European Union or HaDEA, the granting authority. Neither the European Union nor the granting authority can be held responsible for them. We acknowledge the Department of Agriculture, Food and the Marine (DAFM) for making the relevant data publicly available, and recognize the efforts of its staff in collecting, processing, and publishing the data used in this study.

